# *VILMIR* is a *trans*-acting long noncoding RNA that enhances the host interferon response in human epithelial cells

**DOI:** 10.1101/2025.08.18.670784

**Authors:** Kristen John, Ethan Smith, Alexandra Istishin, Nasif Mahmood, Kayleigh Diveley, Tammy S. Tollison, Susan Carpenter, Xinxia Peng

## Abstract

Long noncoding RNAs (lncRNAs) have been found to play significant regulatory roles within antiviral and immune responses. We previously identified the novel lncRNA virus-inducible lncRNA modulator of interferon response (*VILMIR*), that was found to broadly regulate the host transcriptional response to interferon-beta (IFN-β) treatment in A549 human lung epithelial cells. Here, we investigated the mechanism by which *VILMIR* regulates the host interferon response in *trans* by identifying interacting proteins and gene regulatory networks of *VILMIR*. Through an RNA pull-down assay, we found that *VILMIR* interacted with both nuclear and cytoplasmic proteins *in vitro*, including the transcriptional regulators FUBP1 and PUF60 in the nucleus, as well as the antiviral proteins IFIT1 and IFIT3 and the aminoacyl-tRNA synthetases QARS1 and KARS1 in the cytoplasm. In addition, we found that overexpression of *VILMIR* in A549 cells resulted in an overall enhancement of host interferon response genes and identified a core set of interferon-stimulated genes that were consistently regulated by *VILMIR* knockdown and overexpression. Finally, we proposed several possible mechanisms by which *VILMIR* may interact with the identified proteins to regulate the interferon response, such as by interacting with FUBP1 and PUF60 in the nucleus to regulate host transcription in *trans* or by interacting with the IFIT proteins and aminoacyl-tRNA synthetases in the cytoplasm to regulate translation.

**IMPORTANCE:** Despite thousands of long noncoding RNAs (lncRNAs) being differentially expressed after immune responses and viral infections, there is very limited knowledge on their individual functions in these contexts. We previously identified a novel lncRNA, *VILMIR*, that was found to be an interferon-stimulated gene that regulated the host transcriptional response to interferon-beta treatment in human epithelial cells. Here, we investigated the mechanism by which *VILMIR* regulates the interferon response in *trans*. Through *in vitro* studies, we identified several nuclear and cytoplasmic proteins that interact with *VILMIR*, including proteins involved in transcriptional and translational regulation. In addition, we demonstrated that the overexpression of *VILMIR* results in an enhancement of host interferon response genes, supporting our hypothesis that *VILMIR* plays an activating role in the host interferon response. Finally, we propose several potential models for the mechanism of *VILMIR*, providing a foundation for the investigation of *VILMIR* as a novel therapeutic target in antiviral immunity.

## INTRODUCTION

Noncoding RNAs are known to be major regulators of cellular processes, such as ribosomal RNA (rRNA) and transfer RNA (tRNA) which are critical for protein synthesis (1, 2), as well as small nuclear RNA (snRNA) and small nucleolar RNA (snoRNA) which are critical for splicing and RNA modification (3, 4). The largest class of the noncoding RNAs are long noncoding RNAs (lncRNAs), defined as transcripts greater than 500 nucleotides in length with low translational potential (5). The recent GENCODE V47 release estimates 35,934 lncRNA genes in the human genome (6). However, despite the large number of annotated lncRNAs, their functions are still widely unknown.

Individual lncRNAs have been identified to have significant functions within biological processes such as cell development (7), cancer (8), and inflammation (9) by regulating processes such as gene transcription and protein translation (10). There is also growing evidence that lncRNAs play important roles within antiviral and immune responses (11). For example, a recent study in 2023 identified that the overexpression of a lncRNA, LncRNA#61, inhibited influenza A virus (IAV) replication in human cells and even reduced viral replication *in vivo* after lipid nanoparticle-encapsulated delivery of LncRNA#61 in mice (12). As regulatory RNAs such as lncRNAs have low translational potential, they often function by interacting with RNA-binding proteins (RBPs) to regulate transcription (13) or act as scaffolds for protein interactions (14). While many lncRNAs have been found to act as *cis* regulators by regulating the expression of neighboring protein-coding genes (15), lncRNAs have also been found to function as *trans* regulators by regulating transcription on different chromosomes (16). Some lncRNAs can act in both *cis* and *trans*, such as *lincRNA-Cox2* that was found to regulate its neighboring gene, *Ptgs2*, as well as a subset of immune genes in *trans* using mouse models (17). In addition, a single lncRNA can have multiple functions in different cellular compartments, such as PYCARD-AS1 that facilitates DNA methylation at the *PYCARD* promoter in the nucleus, as well as interacts with PYCARD mRNA in the cytoplasm to inhibit ribosome assembly (18). Therefore, understanding the RBP interactions of a lncRNA can help elucidate how it functions within the cell.

We previously identified a novel lncRNA named virus-inducible lncRNA modulator of interferon response (*VILMIR*) that was found to regulate the host transcriptional response to both interferon-beta (IFN-β) treatment and IAV infection in A549 human lung epithelial cells (19). We found that *VILMIR* did not regulate transcription of its neighboring protein-coding genes but rather had a broad transcriptional regulation; however, the exact mechanism of this regulation was not explored. Therefore, in this study, we aimed to identify interacting proteins and gene regulatory networks of *VILMIR* to better understand its molecular interactions and how it regulates the interferon response in *trans*. Using an RNA-pull down assay, we found that *VILMIR* interacts with several proteins in A549 nuclear and cytoplasmic lysate, including the transcriptional regulators FUBP1 and PUF60, as well as antiviral proteins IFIT1 and IFIT3 and aminoacyl-tRNA synthetases QARS1 and KARS1. In addition, we overexpressed *VILMIR* in A549 cells and found that overexpression resulted in an overall enhancement of host interferon response genes, supporting our hypothesis that *VILMIR* plays an activating role in the host interferon response. By combining RNA-seq analyses from both *VILMIR* knockdown and overexpression studies, we further identified a core set of genes that are consistently regulated by *VILMIR* perturbation in *trans*. Finally, we proposed several potential models of *VILMIR* function, suggesting that *VILMIR* may function in both the nucleus and the cytoplasm to regulate host interferon responses.

## MATERIALS AND METHODS

### Cell culture

The human cancer cell line, A549 lung epithelial (CCL-185), was purchased from American Type Culture Collection (ATCC, Manassas, VA, USA). Human embryonic kidney (HEK) epithelial 293FT cells were ordered from Invitrogen (ThermoFisher Scientific). A549 cells were maintained in F-12K media with 10% fetal bovine serum (FBS). HEK 293FT cells were maintained in DMEM media plus 1% glutamax and 10% FBS. All cell lines were kept at 37°C in a 5% CO_2_ incubator and maintained in culture as recommended by ATCC.

### Protein extraction

In order to collect protein lysate for subsequent RNA pull-down, confluent T182 flasks of A549 cells were washed with 1X Dulbecco’s phosphate buffered saline (DPBS) and treated with fresh A549 media containing 1 ng/mL human IFN-β recombinant protein (R&D Systems™ 8499IF010) for six hours. Nuclear and cytoplasmic protein lysate of the cells was extracted using NE-PER Nuclear and Cytoplasmic Extraction Reagents (Thermo Scientific) according to the manufacturer’s instructions. Relative protein quantification was determined using absorbance at 280 nm by comparison with a Bovine Serum Albumin (BSA) standard curve.

### RNA pull-down assay and mass spectrometry

Full-length *VILMIR* (Table S1) or its antisense sequence was cloned into the pGEM-3Z vector (Promega) downstream of the T7 promoter using EcoRI and BamHI restriction enzyme sites. The plasmids were linearized with BamHI on the 3’ end in order to facilitate *in-vitro* transcription. Biotinylated RNA probes were *in-vitro* transcribed by the T7 RNA polymerase using TranscriptAid T7 High Yield Transcription Kit (Thermo Scientific) and a 1:39 molar ratio of Biotin-16-UTP (ApexBio Technology) to standard UTP. Subsequent RNA was purified using the RNeasy Plus Mini Kit (Qiagen) with a genomic DNA eliminator column, and the size was confirmed with an Agilent Bioanalyzer or TapeStation 4150.

The pull-down assay was adapted for RNA from Springer Protocols (20). Briefly, 1.5 mg of prewashed Dynabeads™ M-280 Streptavidin (Invitrogen) was incubated with 25 µg of *VILMIR* sense or antisense RNA probes at room temperature with agitation for 30 minutes. Excess biotinylated RNA probe was removed from the bead-probe complex by several washes. Approximately 3.6 mg of nuclear or 5.3 mg of cytoplasmic protein lysate from A549 cells were incubated with 12.5 µg of antisense bead:probe complexes at 4°C with agitation for 30 minutes as a preclearing step. Precleared lysates were then incubated with *VILMIR* sense or antisense bead:probe complexes with 50 µg tRNA competitor (Thermo Scientific) and 300 U of SUPERase·In™ RNase Inhibitor (Invitrogen) at 4°C with agitation for one hour. Bead:probe:protein complexes were washed three times in washing buffer, eluted in water at 70°C for 5 minutes, and boiled in Laemmli buffer at 95°C for 10 minutes. The pull-down assay was performed in triplicate for both sense and antisense reactions. The supernatant was run on a 6% SDS-PAGE gel for approximately 10 minutes, and then the gel was stained with Coomassie blue and destained to excise and prepare the samples for protein identification by mass spectrometry. Mass spectrometry was performed by the BIDMC-Harvard Mass Spectrometry Facility and Asara Laboratory using the Thermo Scientific QExactive HFx Orbitrap nano HR-LC-MS/MS following in-gel digestion of proteins with Trypsin/LysC.

To identify *VILMIR* sense-specific binding proteins in the nucleus or cytoplasm, we first averaged the peptide spectrum counts in the sense and antisense replicates and then calculated an approximate fold-change (FC) value by taking the difference between the sense and antisense peptide counts (sense/antisense). Proteins were eliminated that were evenly distributed between the sense and antisense samples (i.e. FC of 1), enriched in the antisense samples (FC < 1), or whose replicates had inconsistent counts. *VILMIR* sense-specific bound proteins were identified as those proteins that had protein peptide spectrum counts in all three replicates of the sense RNA but none in the antisense replicates (unique binding), or whose peptide spectrum counts were enriched (FC > 1) in the sense replicates over the antisense replicates (enriched binding). These remaining proteins were narrowed down further by removing proteins with a FC < 2 in the sense replicates, as well as focusing on proteins with known associations with interferon and antiviral responses. The identified proteins can be found in Tables 1 and 2.

**Table 1.** Mass spectrometry identification of *VILMIR*-interacting proteins in A549 nuclear cellular lysate treated with IFN-β as determined by RNA pull-down assay.

| Protein ID | Mass (kDa) | Sense *VILMIR* Peptide Count |  |  | Antisense *VILMIR* Peptide Count |  |  | Enrichment (Avg Sense/Avg Antisense) |
| --- | --- | --- | --- | --- | --- | --- | --- | --- |
| LRPPRC | 158 | 19 | 15 | 15 | 0 | 0 | 0 | N/A |
| PCBP2 | 39 | 10 | 8 | 2 | 0 | 0 | 0 | N/A |
| CSTF3 | 83 | 10 | 9 | 10 | 0 | 0 | 0 | N/A |
| CELF1 | 52 | 11 | 9 | 7 | 0 | 0 | 0 | N/A |
| CSTF2 | 61 | 6 | 5 | 5 | 0 | 0 | 0 | N/A |
| TARDBP | 45 | 6 | 5 | 3 | 0 | 0 | 0 | N/A |
| CPSF7 | 52 | 5 | 4 | 4 | 0 | 0 | 0 | N/A |
| FUBP1 | 68 | 24 | 24 | 23 | 5 | 2 | 2 | 7.9 |
| KHSRP | 73 | 87 | 91 | 84 | 15 | 12 | 9 | 7.3 |
| FUBP3 | 62 | 8 | 7 | 5 | 0 | 0 | 1 | 6.7 |
| ELAVL1 | 36 | 6 | 3 | 3 | 0 | 0 | 2 | 6.0 |
| CSTF1 | 48 | 122 | 133 | 135 | 27 | 26 | 18 | 5.5 |
| SYMPK | 141 | 4 | 7 | 5 | 1 | 0 | 0 | 5.3 |
| SF1 | 68 | 54 | 61 | 65 | 14 | 15 | 14 | 4.2 |
| HNRNPM | 78 | 20 | 23 | 19 | 6 | 5 | 4 | 4.1 |
| PUF60 | 60 | 46 | 55 | 52 | 16 | 18 | 15 | 3.1 |
| CPSF1 | 161 | 29 | 28 | 35 | 10 | 10 | 10 | 3.1 |
| CPSF2 | 88 | 39 | 40 | 38 | 15 | 14 | 13 | 2.8 |
| SCAF11 | 165 | 85 | 89 | 72 | 36 | 37 | 33 | 2.3 |

**Table 2.** Mass spectrometry identification of *VILMIR*-interacting proteins in A549 cytoplasmic cellular lysate treated with IFN-β as determined by RNA pull-down assay.

| Protein ID | Mass (kDa) | Sense *VILMIR* Peptide Count |  |  | Antisense *VILMIR* Peptide Count |  |  | Enrichment (Avg Sense/Avg Antisense) |
| --- | --- | --- | --- | --- | --- | --- | --- | --- |
| IFIT3 | 56 | 4 | 7 | 4 | 0 | 0 | 0 | N/A |
| LARS1 | 134 | 16 | 16 | 9 | 1 | 0 | 0 | 13.7 |
| IFIT1 | 55 | 10 | 12 | 11 | 1 | 1 | 1 | 11.0 |
| QARS1 | 88 | 15 | 14 | 14 | 2 | 1 | 2 | 8.6 |
| AIMP1 | 34 | 8 | 9 | 5 | 1 | 0 | 1 | 7.3 |
| MARS1 | 101 | 17 | 16 | 10 | 2 | 2 | 3 | 6.1 |
| HNRNPK | 51 | 22 | 22 | 17 | 5 | 5 | 3 | 4.7 |
| RARS1 | 75 | 23 | 13 | 15 | 3 | 3 | 5 | 4.6 |
| KARS1 | 68 | 48 | 35 | 45 | 12 | 9 | 9 | 4.3 |
| PCBP2 | 39 | 4 | 3 | 4 | 0 | 1 | 0 | 3.7 |
| FUBP1 | 68 | 31 | 38 | 37 | 12 | 10 | 11 | 3.2 |
| IARS1 | 145 | 36 | 40 | 34 | 13 | 10 | 13 | 3.1 |
| SART3 | 110 | 12 | 11 | 7 | 4 | 3 | 3 | 3.0 |
| PUF60 | 60 | 16 | 22 | 20 | 7 | 6 | 10 | 2.5 |
| U2AF2 | 54 | 13 | 11 | 11 | 6 | 6 | 2 | 2.5 |
| DARS1 | 57 | 21 | 19 | 20 | 9 | 7 | 10 | 2.3 |
| LRPPRC | 158 | 138 | 141 | 141 | 68 | 62 | 55 | 2.3 |
| EPRS1 | 171 | 35 | 35 | 18 | 12 | 14 | 13 | 2.3 |
| SF3B3 | 136 | 8 | 11 | 7 | 3 | 4 | 5 | 2.2 |

### Western blotting

Identified proteins from mass spectrometry were confirmed by Western blot after a pull-down assay as described above. Eluted protein from the *VILMIR* pull-down assay was loaded into 10% SDS-PAGE gels and transferred to polyvinylidene difluoride (PVDF) membranes (Invitrogen). Membranes were blocked for either 1 hour at room temperature or 4°C overnight in 1X Tris Buffered Saline (TBS) with 1% (w/v) Casein and then incubated in primary antibody diluted in blocking buffer for 1 hour at room temperature. The following primary antibodies were used: anti-FUBP1 1:1000 (Proteintech, 24864-1-AP), anti-PUF60 1:1000 (Proteintech 10810-1-AP), anti-PCNA 1:5000 (Proteintech, 10205-2-AP), anti-IFIT1 1:500 (Cell Signaling Technology, #14769), anti-IFIT3 1:2000 (Proteintech, 15201-1-AP), anti-GlnRS (or QARS1) 1:2000 (Proteintech 12645-1-AP), anti-KARS 1:1000 (Proteintech 14951-1-AP) and anti-GAPDH 1:5000 (Proteintech, 10494-1-AP). Membranes were rinsed 2X and washed 2X for 5 minutes each in TBS buffer with 0.05% Tween 20 (TBS-T) and then incubated in Goat anti-Rabbit IgG (H+L) secondary antibody, HRP conjugate (Invitrogen #31460) diluted 1:5,000 in blocking buffer for 30 minutes at room temperature. Membranes were rinsed 3X and washed 3X for 5 minutes each in TBS-T buffer and then detected by chemiluminescence using either Pierce™ ECL Substrate (Thermo Scientific) for the FUBP1 and PCNA blots (Fig 1A) or SuperSignal™ West Atto Ultimate Sensitivity Substrate (Thermo Scientific) for the remaining blots (PUF60 in Fig 1A and B-C), according to the detection limits of the protein. The blots were visualized using a Bio-Rad ChemiDoc MP Imaging System. Densitometry analysis was performed using ImageJ software where applicable. Briefly, the background was substracted and the density of each band was measured. The sense and antisense bands were then normalized to their input band and the average fold change of three independent replicates was calculated. FUBP1 and PUF60 in Fig 1A were not included in the densitometry analysis as the antisense band was too low to obtain an accurate density.

**Figure 1.**
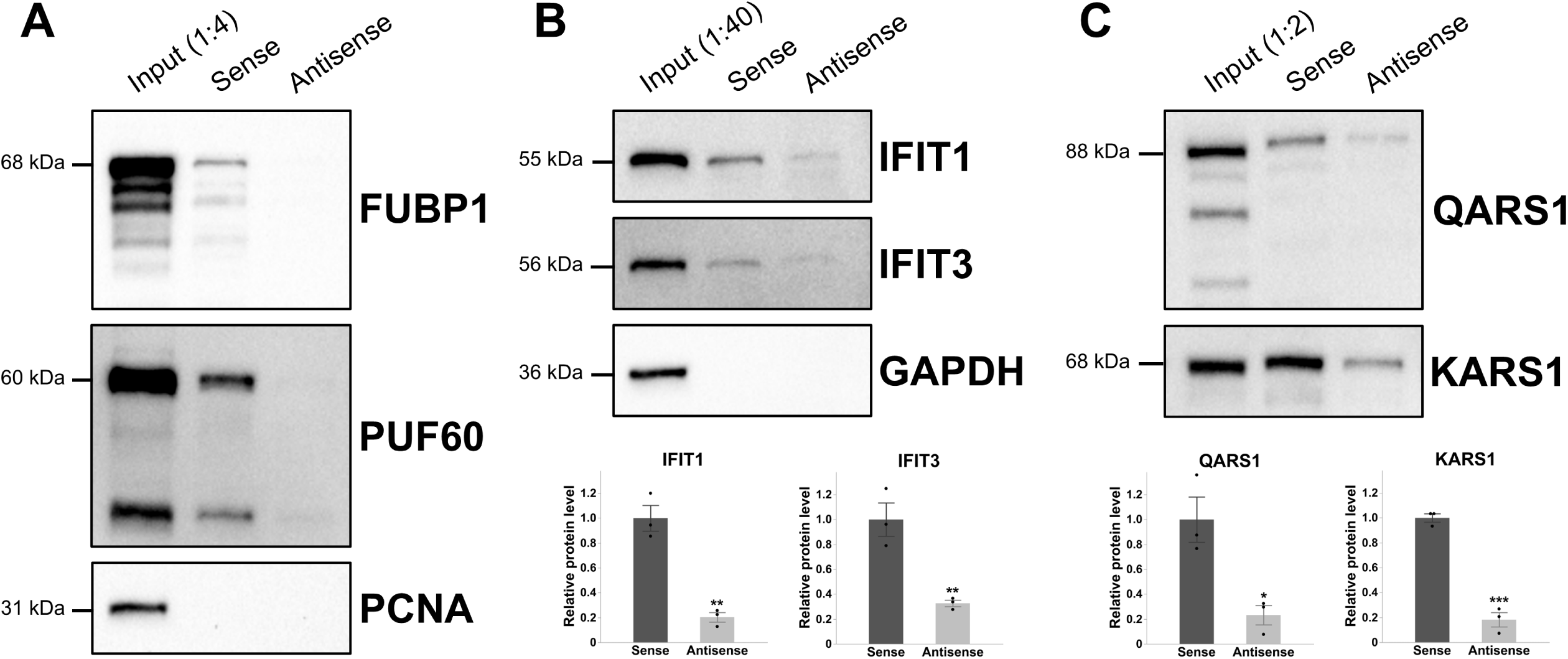
*VILMIR* sense RNA interacts with nuclear and cytoplasmic proteins from A549 lysate. An RNA pull-down assay was performed by incubating *in-vitro* transcribed *VILMIR* sense RNA or antisense RNA (negative control) with nuclear or cytoplasmic lysate from IFN-treated A549 cells. Interacting proteins were identified by mass spectrometry. The full list of identified proteins can be found in Tables 1 and 2. A) The interaction of *VILMIR* sense RNA with FUBP1 and PUF60 was confirmed by Western Blot. PCNA was used as a negative nuclear control, and input protein was diluted 1:4. B) The interaction of *VILMIR* sense RNA with IFIT1 and IFIT3 was confirmed by Western blot with GAPDH as a negative cytoplasmic control, and input protein was diluted 1:40. Densitometry analysis of IFIT1 and IFIT3 protein bands in the Western blots are displayed below the blots. C) The interaction of *VILMIR* sense RNA with QARS1 and KARS1 was confirmed by Western blot, and input protein was diluted 1:2. Densitometry analysis of QARS1 and KARS1 protein bands in the Western blots are displayed below the blots. All Western blots are representative of three independent replicates. Proteins were detected by chemiluminescence using either Pierce™ ECL Substrate (Thermo Scientific) for the FUBP1 and PCNA blots (A) or SuperSignal™ West Atto Ultimate Sensitivity Substrate (Thermo Scientific) for the remaining blots (PUF60 in A and B-C). Densitometry analysis was not performed for panel A as the antisense band was too low to obtain an accurate density. *P < 0.05, **P < 0.01, ***P < 0.001 (Student’s t-test).

### Interferon treatment

Following the same methods as (19), A549 cells were seeded overnight between 150,000-175,000 cells per well in 12-well plates in 1.5 mL media. The following day, the cell monolayer was washed with 1X DBS and treated with fresh A549 media with or without human IFN-β recombinant protein at the indicated concentrations. All cells were harvested at six hours after treatment according to the TRIzol Reagent User Guide (Invitrogen).

### RNA isolation and quantitative PCR

Total RNA was isolated from cells following the TRIzol isolation method (Invitrogen) and quantified using Nanodrop spectrophotometry. One μg of RNA was reverse-transcribed into cDNA using the QuantiTect Reverse Transcription Kit (Qiagen) containing both oligo-dT and random primers. Quantitative PCR (qPCR) was performed on the cDNA using PowerUp SYBR Green Master Mix (Applied Biosystems). Relative expression of the indicated RNAs was determined using the ΔΔCt method with GAPDH as an endogenous control. Statistical analysis of significance was performed in JMP Pro 16 software (SAS Institute Inc., Cary, NC). The primer sequences used in this study are as follows: *GAPDH* F: GGTATCGTGGAAGGACTCATGAC; *GAPDH* R:

ATGCCAGTGAGCTTCCCGTTCAG (21); *VILMIR* F: GCTCCACCCTGAAAGTC; *VILMIR* R: CTACACAGTGCTGAGGAAA (19).

### Plasmid Construction and Overexpression of *VILMIR*

The pSico bidirectional expression vector was a gift from Susan Carpenter and described in (22). Full-length *VILMIR* was cloned into the pSico vector with an EF1a promoter expressing zeocin resistance and GFP as a selection marker, using PspXI and NotI restriction enzyme sites. The sequence was confirmed by Sanger sequencing. To produce lentivirus, HEK-293FT cells were co-transfected with the pSico vector expressing *VILMIR* and lentiviral vectors, psPAX2 (Addgene #12260) and pMD2.G (Addgene #12259) using the Lipofectamine 3000 Reagent (Invitrogen). In addition, an empty pSico vector was transfected as a negative control. Viral supernatant was collected 72 hours post-transfection and filtered through a 0.22-µM syringe filter. A549 cells were transduced with lentivirus, and stable integrants were sorted based on GFP expression at the UNC Flow Cytometry Core Facility (Chapel Hill, North Carolina) using a Becton Dickinson FACSAria II. The successful overexpression of *VILMIR* was confirmed by RT-qPCR.

### cDNA library construction, RNA-sequencing, and Ingenuity Pathway Analysis (IPA)

mRNA sequencing was performed in biological triplicate in A549 *VILMIR*-overexpressing and control cell lines treated with mock or either 1 ng/mL or 10 ng/mL human IFN-β for 6 hours. Total RNA was isolated from cells following the TRIzol isolation method (Invitrogen). All samples were quantified and assayed to confirm a minimum RNA integrity number of at least 9.7 using an Agilent TapeStation 4150. Next, 500 ng of total RNA per sample underwent mRNA capture and was then fragmented at 94°C for 6 min. Sequencing libraries were prepared according to the manufacturer’s protocol using 11 cycles of final amplification (KAPA mRNA HyperPrep Kit, catalog no. KK8580 and KAPA UDI Adapter Kit, catalog no. KK8727). Libraries underwent QC prior to sequencing using an Agilent TapeStation 4150. Next-generation sequencing was performed on a Complete Genomics DNBSEQ-G400C (150 bp paired end) to a targeted depth of ∼20 million reads per sample. The sequencing data from *VILMIR* knockdown in (19) was also incorporated.

Complete Genomics RNA-seq reads were mapped against the Hg38 using STAR version 2.7.9 a (23). Custom STAR parameters were set as follows: limitOutSAMoneReadBytes: 1,000,000, outSAMprimaryFlag: AllBestScore, outFilterType: BySJout, alignSJoverhangMin: 8, alignSJDBoverhangMin: 3, outFilterMismatchNmax: 999, alignIntronMin: 20, alignIntronMax: 1,000,000, alignMatesGapMax: 1,000,000, outFilterMultimapNmax: 20; otherwise, default STAR parameters were used. Following read mapping, a count matrix was generated from the STAR results using R. Genes were removed from the matrix if they did not have at least 30 reads in a minimum of three samples from either the perturbation group (knockdown or overexpression) or the control group. Counts were normalized using the TMM normalization method via the calcNormFactors function in edgeR version 3.40.2 (24).

To conduct our differential gene expression analysis, we utilized the limma-trend approach from Limma version 3.54.2 (25). For each dose level of the IFN-β treatment, we assessed differential expression by comparing each knockdown or overexpression condition to its respective mock treatment. We then contrasted the differential expression results of each knockdown or overexpression (STAT1g1/1 ng IFN-β vs STAT1g1/Mock, VILMIRg1/1 ng IFN-β vs VILMIRg1/Mock, etc.) against that of its corresponding control (Ctrl/1 ng IFN-β vs Ctrl/Mock, Ctrl/10 ng IFN-β vs Ctrl/Mock). Genes were considered differentially expressed in a given contrast if their unadjusted P-value was less than 0.05 with no fold change requirement. To identify IFN-responsive genes, we further filtered these results by requiring a log2FC greater than 1.25 following IFN-β treatment in the control cell lines for each experiment. Differential expression results were visualized using the ComplexHeatmap R package version 2.14.0 (26).

Pathway enrichment analysis was generated using QIAGEN Ingenuity Pathway Analysis (IPA) (27). A raw *P*-value cutoff of < 0.05 was used to define genes with significant expression changes after *VILMIR* overexpression in each IFN-β treatment. Canonical pathways analysis identified the pathways from the QIAGEN IPA library of canonical pathways that were most significant to the data set. Differentially expressed genes from the data set that met the *P*-value cutoff of 0.05 (−log10 *P*-value 1.3) and were associated with a canonical pathway in the QIAGEN Knowledge Base were considered for the pathway analysis. A right-tailed Fisher’s exact test was used to calculate a *P*-value determining the probability that the association between the genes in the data set and the canonical pathways is explained by chance alone.

## RESULTS

### LncRNA *VILMIR* can interact with nuclear and cytoplasmic proteins from A549 epithelial cells *in vitro*, including FUBP1, PUF60, IFIT1, IFIT3, QARS1, and KARS1

To identify potential protein interactions of *VILMIR*, we performed an RNA pull-down assay. As *VILMIR* was localized in both nuclear and cytoplasmic compartments in A549 human lung epithelial cells (19), we investigated its potential protein interactions in both compartments*. In vitro-*transcribed biotinylated *VILMIR* as well as antisense *VILMIR* control RNA was incubated with nuclear or cytoplasmic lysates of A549 cells treated with IFN-β to mimic cellular interactions during a host interferon response. Interacting RBPs were then identified by mass spectrometry. The full list of identified proteins was narrowed down by prioritizing proteins that either uniquely interacted with *VILMIR* sense RNA compared to antisense RNA or were greater than two times enriched in the *VILMIR* sense RNA compared to antisense RNA according to peptide spectrum counts (see Materials and Methods). From these criteria, we identified 19 proteins in each compartment that were either unique to or enriched in the *VILMIR* sense RNA (Tables 1 and 2). This suggests that *VILMIR* may have functions in both compartments.

In nuclear A549 lysate, *VILMIR* sense RNA was found to interact with several proteins involved in pre-mRNA splicing and processing (CSTF1, CSTF2, CSTF3, CELF1, CPSF7, SYMPK, SF1, HNRNPM, CPSF1, CPSF2, and SCAF11), as well as transcriptional regulation (TARDBP, FUBP1, KHSRP, FUBP3, and PUF60). Using Western blot analysis, we confirmed the interaction of *VILMIR* sense RNA with FUBP1 and PUF60 (Figure 1A). FUBP1, or Far Upstream Element Binding Protein 1, acts as a transcriptional regulator and is a well-known activator of the *c-Myc* oncogene (28). While FUBP1 is primarily located in the nucleus, it has been found to translocate to the cytoplasm (29) and was also identified in our cytoplasmic mass spectrometry results (Table 2). To negatively control expression of *c-Myc*, FUBP1 also interacts with the FUBP-Interacting Repressor (FIR), which is an alternatively spliced variant of Poly(U) Binding Splicing Factor 60 (PUF60), meaning that FUBP1 can be involved in both positive and negative regulation of gene expression (28).

Interestingly, in cytoplasmic A549 lysate, *VILMIR* sense RNA interacted with IFIT1 and IFIT3 proteins more than the antisense RNA (Table 2). The IFIT protein family, or IFN-induced protein with tetratricopeptide repeats, are IFN-stimulated genes (ISGs) that get induced during antiviral immune responses and consist of *IFIT1*, *IFIT2*, *IFIT3*, and *IFIT5* in humans (30). The IFIT proteins are well-known to inhibit translation of both cellular mRNA and viral RNA by either interacting with eukaryotic initiation factor 3 (eIF3) to block translation (31) or by binding directly to the 5’ end of non-self RNAs (32, 33). While IFIT2 and IFIT5 were not identified as *VILMIR*-interacting proteins in the mass spectrometry, the interaction of *VILMIR* sense RNA with IFIT1 and IFIT3 proteins was confirmed by Western blot and densitometry analysis (Figure 1B). In addition, *VILMIR* sense RNA was found to interact with eight aminoacyl-tRNA synthetases (ARSs), which are enzymes responsible for pairing tRNAs with amino acids during translation, as well as ARS-interacting multifunctional protein 1 (AIMP1), which helps form the multi-tRNA synthetase complex (34) (Table 2). Two of these ARSs were confirmed by Western blot and densitometry analysis, QARS1 or glutaminyl-tRNA synthetase 1, and KARS1 or lysyl-tRNA synthetase 1 (Figure 1C). Therefore, while several nuclear and cytoplasmic protein interactions were confirmed *in vitro* and should be confirmed in the cells, these results suggest that *VILMIR* could function in the nucleus by interacting with proteins such as FUBP1 and PUF60 to regulate transcription or in the cytoplasm by interacting with IFIT1 and IFIT3 and/or QARS1 and KARS1 to regulate translation.

### *VILMIR* overexpression results in minimal fold change differences before interferon-β treatment in A549 epithelial cells

As our pull-down assay suggested that *VILMIR* could function through the transcript itself, we were interested if overexpression of *VILMIR* in cells would cause significant expression changes, both before and after IFN-β treatment. Therefore, we generated an A549 cell line overexpressing ectopic *VILMIR* and performed RNA-sequencing (RNA-seq) analysis of overexpressing cells treated with or without two separate concentrations of human IFN-β. Compared to the control cell line with an empty vector, we observed a 7.5-fold increase in the baseline expression of *VILMIR* (Figure 2A). We first examined if *VILMIR* alone caused significant expression differences outside of an IFN response by comparing gene expression in the mock-treated cell lines. With a relaxed criterion for differential expression analysis (raw *P*-value < 0.05 and no fold change cutoff), we identified 459 genes that showed altered expression changes in the *VILMIR*-overexpressing cell line compared to the control cell line in the mock-treatment (Figure 2B).

**Figure 2.**
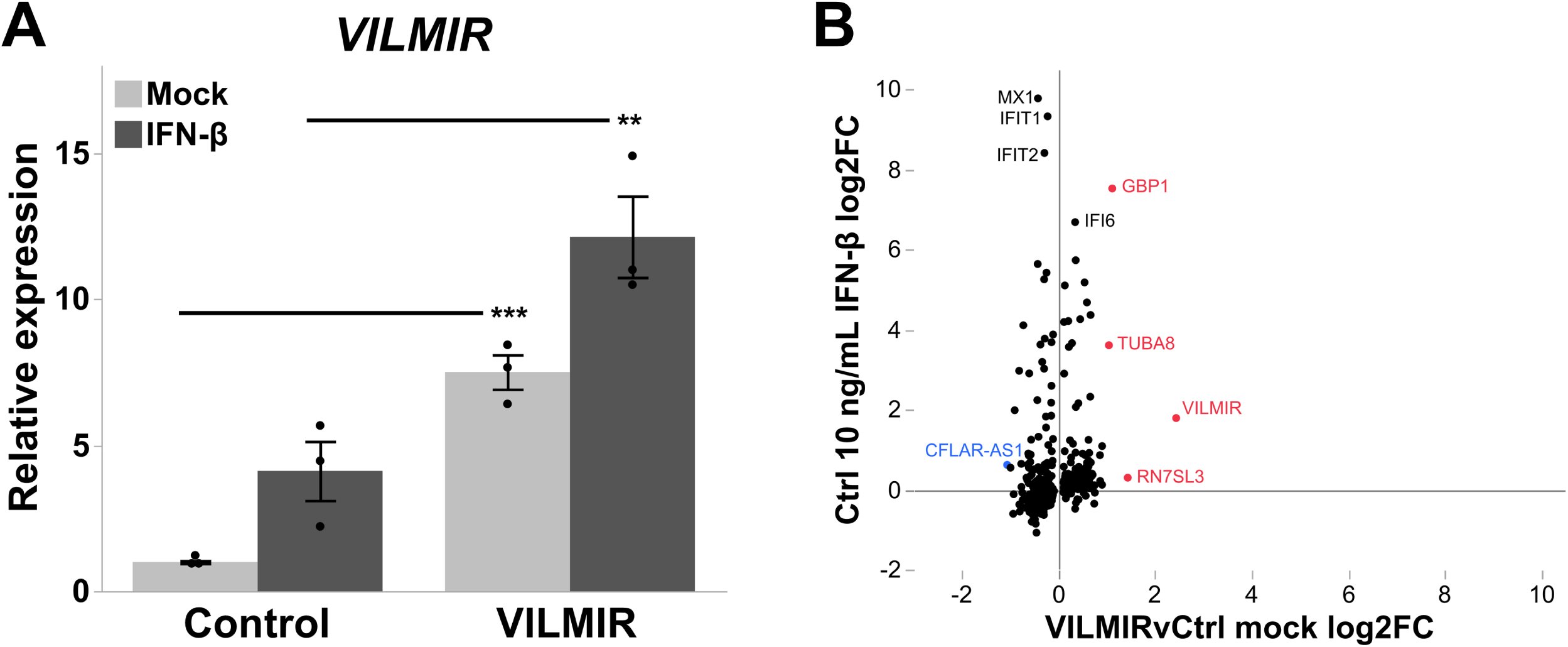
Overexpression of *VILMIR*. (A) A549 cells were transduced with a vector expressing ectopic *VILMIR* or an empty-vector control and treated with mock or 10 ng/mL human IFN-β for 6 hours. Relative expression of *VILMIR* was determined by RT-qPCR and normalized to the mean of the mock-treated control cell line. Data was normalized to GAPDH using the ΔΔCt method and expressed as means ± SE (n=3). **P < 0.01, ***P < 0.001 (Student’s t-test). (B) RNA-seq analysis was performed to identify genes that showed altered expression changes after *VILMIR* overexpression in the mock and IFN-treated cells. Displayed are points representing 459 genes that exhibited significant expression changes in the *VILMIR*-overexpressing cell line compared to the control cell line in the mock-treated cells (raw P-value < 0.05). The x-axis indicates the log2FC difference between each gene in the *VILMIR*-overexpressing cells versus the control cells (VILMIRvCtrl) in the mock, while the y-axis represents the log2FC of those same genes after 10 ng/mL IFN-β treatment in the control cell line. The five most differentially expressed genes are labeled in either red (upregulation) or blue (downregulation), as well as four other interferon-stimulated genes of interest in black.

Interestingly, there were several ISGs impacted by *VILMIR* overexpression in the mock-treated cells, including *MX1*, *IFIT1*, *IFIT2*, and *IFI6*, suggesting that *VILMIR* expression may regulate ISGs before IFN-treatment. However, when plotting these same genes against their log2 fold change (log2FC) after IFN-β treatment in the control cell line, we observed that the log2FC differences between the mock-treated cell lines were relatively small in comparison. In fact, only 14% of the total 459 genes exhibited absolute fold changes greater than 1.5, and five genes exhibited an absolute fold change greater than 2 (Figure 2B). Apart from *VILMIR* itself, which was the highest upregulated gene as expected, *RNS7SL3*, *GBP1*, and *TUBA8* were upregulated and *CFLAR-AS1* was downregulated with a fold change greater than 2 (Figure 2B). As *VILMIR* was transcribed ectopically from an overexpression vector, this suggests that *VILMIR* can function through its transcript, rather than just transcription at its genomic locus. In addition, these results suggest that *VILMIR* may have a regulatory role outside of the IFN response, particularly with transcription of *RNS7SL3*, *GBP1*, *TUBA8*, and *CFLAR-AS1*. However, as the majority of expression differences caused by *VILMIR* in the mock-treated cells were relatively small, we sought to determine the impact of *VILMIR* overexpression on host transcription in response to IFN-β treatment.

### Overexpression of *VILMIR* enhances the host transcriptional response to interferon-β treatment in A549 epithelial cells

Next, we determined the impact of *VILMIR* overexpression on the host transcriptional response to IFN-β treatment using the same RNA-seq analysis described above. To investigate the potentially broad regulatory roles of *VILMIR*, similarly as in our previous study (19), we first used a relaxed criterion for differential expression analysis, i.e., raw P-value < 0.05. Using this criterion, we identified 731 genes that showed altered expression changes to IFN-β treatment after *VILMIR* overexpression in at least one of the two doses of IFN-β (Table S2). When analyzing the host transcriptional response to IFN-β, we observed larger fold change differences after *VILMIR* overexpression, with 33-35% of differentially expressed genes (DEGs) exhibiting absolute fold changes greater than 1.5 in either IFN-β treatment, compared to 14% in the mock treatment, indicating that *VILMIR* overexpression has larger impacts during an IFN response.

To identify canonical pathways enriched in the DEGs impacted by *VILMIR* overexpression, QIAGEN Ingenuity Pathway Analysis (IPA) was performed (27). There were 17 canonical pathways significantly enriched in the overexpression cell line in both IFN-β treatments with a raw enrichment

P-value < 0.05 (−log10 P-value >1.3), with the most significant of these pathways being the Interferon alpha/beta signaling pathway (Figure 3A and Table S3). In addition, IPA predicted an overall activation of this pathway, with positive z-scores of 2.121 and 1 for the 1 ng/mL and 10 ng/mL IFN-β treatments, respectively. The majority of the genes represented in this pathway such as *MX1*, *IFIT2*, *IRF7*, *IFIT1*, *RNASEL*, *TYK2*, and *OAS1* had higher fold changes after *VILMIR* overexpression compared to the control cell line, besides *IFITM1* which had a lower fold change (Figure 3B). However, it is possible *IFITM1* could serve as a negative regulator, as it has previously been found to be a negative regulator in certain contexts, with suppression of *IFITM1* inhibiting proliferation in glioma cells (35). The overall enhancement of genes within the IFN pathway after *VILMIR* overexpression is consistent with our previous findings that *VILMIR* knockdown dampened ISGs (19), supporting our hypothesis that *VILMIR* plays an activating role in the host interferon response.

**Figure 3.**
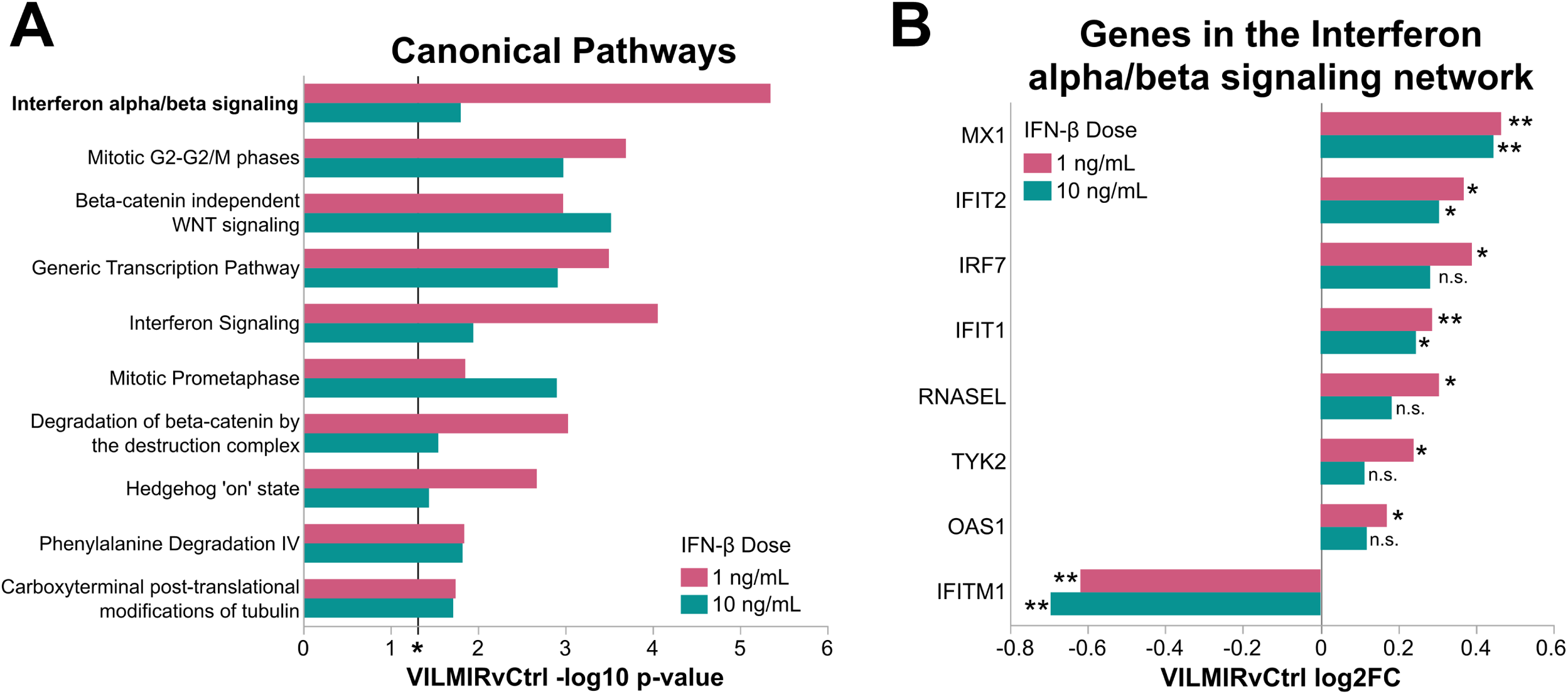
Overexpression of *VILMIR* results in the enhancement of interferon-stimulated genes after IFN-β treatment in A549 cells. (A) RNA-seq analysis was performed to identify genes that showed altered expression changes to IFN-β treatment after *VILMIR* overexpression in at least one of two doses of IFN-β (raw P-value < 0.05). QIAGEN IPA was then performed to identify canonical pathways significantly enriched in the DEGs impacted by *VILMIR* overexpression after 1 ng/mL or 10 ng/mL IFN-β. Shown are the top 10 out of 17 significant pathways shared between both IFN-β treatments (Table S2). Enriched pathways that met the raw enrichment P-value < 0.05 (−log10 P-value cutoff of 1.3) using a right-tailed Fisher’s exact test and were associated with a canonical pathway in the QIAGEN Knowledge Base were included here (*P < 0.05 included as reference). (B) Displayed are the genes in the interferon alpha/beta signaling network according to IPA, along with their log2FC values after *VILMIR* overexpression compared to the control (VILMRvCtrl). *P < 0.05, **P < 0.01.

### *VILMIR* knockdown and overexpression consistently regulates the transcription of a core set of interferon-stimulated genes in A549 epithelial cells

In order to identify a core set of genes that were differentially expressed in response to IFN-β treatment after both *VILMIR* overexpression and knockdown, we next combined the RNA-seq analysis of *VILMIR* overexpression with our previous analysis of *VILMIR* knockdown (19), which included two A549 *VILMIR* knockdown (KD) cell lines (VILMIRg1 and VILMIRg2) as well as a *STAT1* KD cell line (STAT1g1) as a positive control for interferon response, treated with the same two IFN-β doses. To obtain a more robust set of genes that are IFN-responsive, we applied an additional filtering step of a fold change greater than 1.25 after IFN-β treatment in the control cell lines for each experiment. After applying this criterion, we then examined expression changes after *VILMIR* KD or overexpression with a raw P-value < 0.05. This gave us a list of 132 IFN-responsive genes that showed altered expression changes to IFN-β treatment after either *VILMIR* KD or overexpression, in at least one of the two IFN-β doses (Figure S1). Out of these genes, 96 were only differentially expressed after *VILMIR* KD, whereas 25 genes were only differentially expressed after *VILMIR* overexpression. This difference in the number of DEGs may be because the KD study contained two *VILMIR* KD cell lines, whereas the overexpression experiment only had a single *VILMIR* overexpression cell line. Another reason could be due to the difference in the method of gene perturbation, as the knockdown targeted endogenous *VILMIR*, whereas the overexpression produced ectopic *VILMIR* transcripts lacking modifications. The remaining 11 out of 132 genes impacted by *VILMIR* perturbation were differentially expressed after both *VILMIR* KD and overexpression, which is what we chose to focus on (Figure 4). One of these genes included *VILMIR* itself, which was expected.

**Figure 4.**
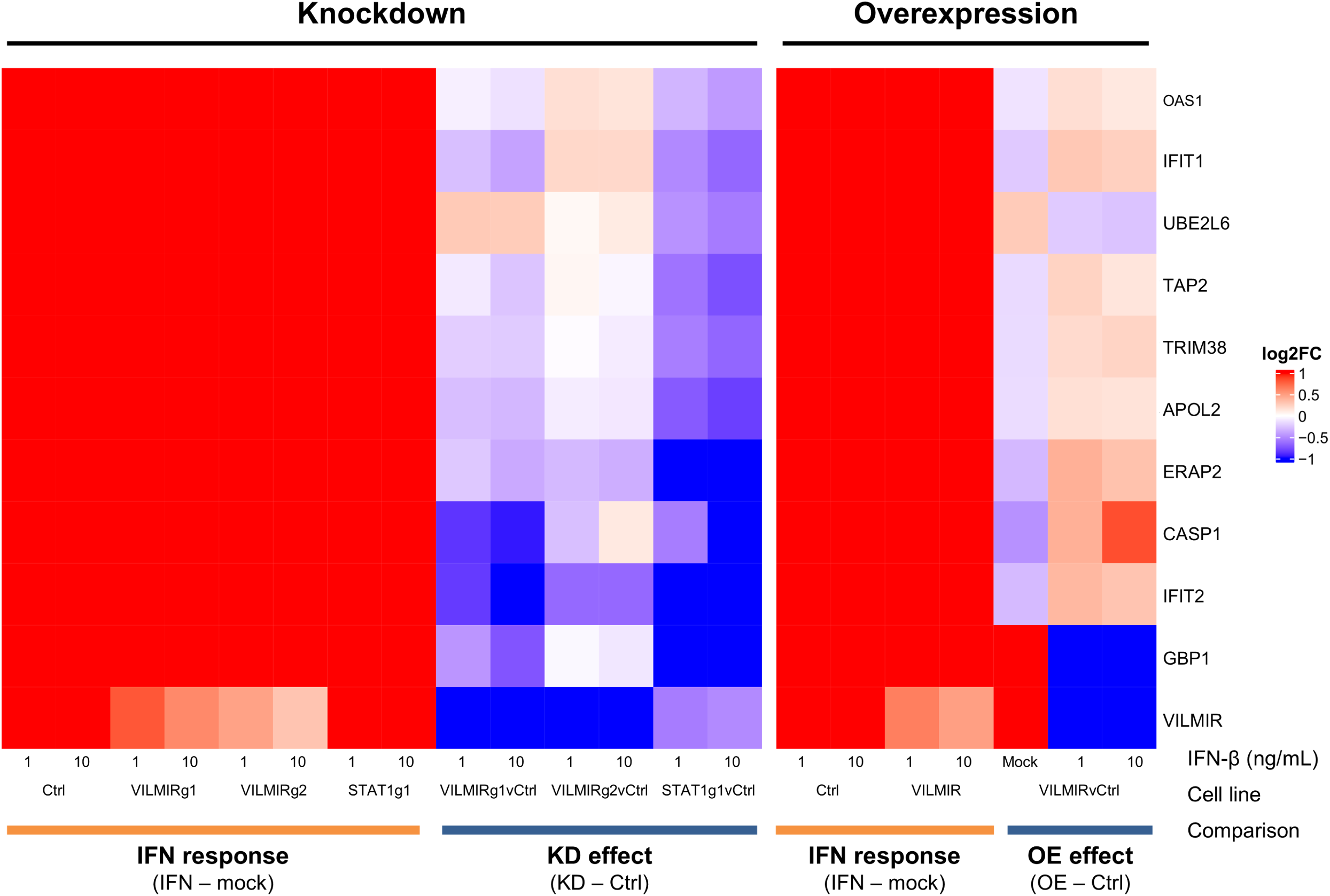
Subset of interferon-stimulated genes that are differentially expressed after *VILMIR* knockdown and overexpression in A549 cells. Heatmap overview of the RNA-seq analysis of two independent experiments: *VILMIR* knockdown (VILMIRg1 and VILMIRg2), *STAT1* knockdown (STAT1g1), or control (Ctrl) A549 gRNA cell lines; and *VILMIR* overexpression (VILMIR) or control (Ctrl) A549 cell lines; all of which were treated with mock or either 1 ng/mL or 10 ng/mL human IFN-β for 6 hours (*n* = 3). The heatmap displays 11 human genes that exhibited significant changes in their responses to IFN-β treatment after both *VILMIR* KD and overexpression (OE), in at least one of two doses of IFN (raw *P*-value <0.05). Rows are genes and columns are conditions and comparisons. As shown by the labels at the bottom, the log2FC after IFN-β treatment in each cell line was first calculated (“IFN response”), and then the “KD effect” or “OE effect” was calculated by comparing the “IFN response” log2FC of each KD/OE line to the “IFN response” log2FC of the control cell line. Red color indicates positive log2FC value (i.e., upregulation) in columns above the label “IFN response,” or higher log2FC values in KD/OE cells compared to that of control cells in columns above the label “KD/OE effect.” The blue color indicates lower log2FC values in KD/OE cells compared to that of control cells in columns above the label “KD/OE effect.”

The 10 genes that were consistently impacted by *VILMIR* perturbation consisted of *OAS1*, *IFIT1*, *UBE2L6*, *TAP2*, *TRIM38*, *APOL2*, *ERAP2*, *CASP1*, *IFIT2*, and *GBP1*, which have all been associated with interferon and antiviral responses in literature (Table 3). As expected, these genes were all upregulated after IFN-β treatment in our A549 cell lines. However, after *VILMIR* knockdown, we observed an overall suppression of the expression of these genes, whereas after *VILMIR* overexpression, these same genes showed an increase in expression (Figure 4). The opposite trend was true for *UBE2L6*, which showed an increase in the *VILMIR* knockdown lines and a decrease in the *VILMIR* overexpression line. However, one reason for this opposite trend may be that *UBE2L6*, a ubiquitin conjugating enzyme, can be a negative regulator in certain contexts, such as inhibiting autophagy in cancer cells (36). Additionally, *GBP1* was decreased in both *VILMIR* knockdown and overexpression after IFN-β treatment. However, as *GBP1* was already upregulated in response to *VILMIR* overexpression before IFN-β treatment, it is possible that *VILMIR* upregulates *GBP1* independently of IFN responses (Figure 4, “Overexpression” mock column, as well as in Figure 2B), which explains why the addition of IFN in the overexpression cell line results in a smaller fold change. This is also true of *VILMIR* itself, which shows a negative fold change difference in the IFN-treated overexpression cells. However, because *VILMIR* is already highly upregulated before IFN-β treatment in the overexpression cells, the addition of IFN-β does not result in a higher fold change compared to the control cells. Finally, as all 10 genes consistently regulated by *VILMIR* are located on different chromosomes than *VILMIR*, these results suggest that *VILMIR* is a *trans* regulator of gene expression.

**Table 3.** Summary of DEGs impacted by both *VILMIR* knockdown and overexpression as seen in Figure 4.

| Gene | Function | Literature |
| --- | --- | --- |
| *OAS1* | Oligoadenylate synthetase 1 – recognizes viral dsRNA and activates RNase L degradation and inhibit translation | (51) |
| *IFIT1* | Binds with eukaryotic initiation factor 3 (eIF3) to block translation; also binds directly to non-self RNAs to inhibit their translation | (31, 33) |
| *UBE2L6* | Ubiquitin conjugating enzyme E2 L6; role in protein degradation and ISGylation | (52) |
| *TAP2* | Transporter associating with antigen processing; transports cytosolic peptides into the ER for presentation by MHC class I molecules | (53) |
| *TRIM38* | E3 ubiquitin ligase; negatively regulates TLR3-mediated type I IFN signaling by targeting TRIF for degradation | (54) |
| *APOL2* | Involved in lipid transport and metabolism; also involved in maintaining airway epithelial layer integrity | (55) |
| *ERAP2* | ER-localized aminopeptidase that plays a role in antigenic peptide editing quality control | (56) |
| *CASP1* | Cysteine-aspartic acid protease; proteolytically cleaves and activates the pro-inflammatory cytokines, IL-1β and IL-18 | (57) |
| *IFIT2* | Binds with eukaryotic initiation factor 3 (eIF3) to block translation; also binds directly to non-self RNAs to inhibit their translation | (31, 33) |
| *GBP1* | GTPase guanylate binding protein; forms an antimicrobial coat to capture cytosol-invasive bacteria and pathogen-containing vacuoles | (58) |

## DISCUSSION AND PROPOSED MODELS OF *VILMIR* FUNCTION

We previously identified the human lncRNA *VILMIR* as a novel ISG during viral infection and found that KD of *VILMIR* in A549 cells resulted in a suppression of the host transcriptional response to IFN-β treatment and IAV infection. However, the mechanism by which *VILMIR* regulates the host interferon response was not explored. Therefore, in this study, we aimed to identify potential protein interactions as well as gene regulatory networks of *VILMIR* to better understand its molecular interactions and propose models of how it may function during an interferon response.

Using an RNA pull-down assay and mass spectrometry, we found that *VILMIR* RNA interacted with several proteins in nuclear and cytoplasmic lysate from A549 epithelial cells treated with IFN-β. As we previously determined that *VILMIR* is distributed in both the nucleus and the cytoplasm (19), these new results further support that *VILMIR* may function in both compartments through protein interactions. Several lncRNAs have also been found to have dual functions in the nucleus and cytoplasm (18, 37–39). For example, lncRNA HOTAIR can function in the nucleus to regulate gene expression by interacting with histone methyltransferases (37), whereas in the cytoplasm it can act as a competing endogenous RNA (ceRNA) by interacting with microRNAs (miRNAs) and regulating translation (38). Therefore, it is possible that *VILMIR* could function through different mechanisms in each compartment as well.

Since KD of *VILMIR* in A549 cells resulted in a suppression of the host transcriptional response to IFN-β treatment and IAV infection, we predicted that *VILMIR* may play an activating role in the host IFN-β response (19). Here, we analyzed the impact of *VILMIR* overexpression during IFN-β treatment. Using RNA-seq analysis, we identified 731 genes that showed altered expression changes to IFN-β treatment after *VILMIR* overexpression in at least one of the two doses of IFN-β. These DEGs were enriched for the Interferon alpha/beta signaling pathway and displayed an overall enhancement of several ISGs after *VILMIR* overexpression, strongly supporting our hypothesis that *VILMIR* plays an activating role in the host IFN response.

By combining our *VILMIR* knockdown and overexpression RNA-seq analysis, we obtained a list of 10 IFN-responsive genes with significant expression changes after both *VILMIR* knockdown and overexpression. Interestingly, seven of the ten genes have known functions related to translational regulation, post-translational regulation by ubiquitination, or protease activity - *OAS1*, *IFIT1*, *UBE2L6*, *TRIM38*, *ERAP2*, *CASP1*, and *IFIT2*. This may mean that *VILMIR* has a regulatory role within translational control or protein processing, which are important cellular processes during a viral infection, as the host translation is tightly regulated in order to limit viral propagation (40). These results were also interesting given our pull-down assay that confirmed the interaction of *VILMIR* with several proteins involved in translation regulation *in vitro*, such as IFIT1, IFIT3, QARS1 and KARS1. Therefore, it is possible *VILMIR* could be regulating ISG expression in the nucleus to modulate their protein abundances, as well as interacting with translational machinery in the cytoplasm. Future work is necessary to determine the potential impact of *VILMIR* on global translation, such as by proteomic analysis or ribosome profiling (41).

Taking these results together, we suggest several potential models for the function of *VILMIR* that should be further explored. First, as *VILMIR* was found to interact with FUBP1 and PUF60 *in vitro*, known transcriptional regulators in the nucleus (28), we suggest a mechanism by which *VILMIR* interacts with proteins such as FUBP1 and PUF60 to regulate gene transcription in *trans* (Figure 5A), as we also observed that *VILMIR* knockdown and overexpression impacts expression of genes through RNA-seq analysis. FUBP1 has been found to be both a positive regulator of transcription as well as a negative regulator through interacting with the repressor protein FIR or PUF60 (28) and has also been associated with virus infection (29, 42). A different lncRNA, NR-109, was previously found to interact with FUBP1 by preventing ubiquitin-mediated degradation of FUBP1 and thus activating *c-Myc* transcription (43). Therefore, as *VILMIR* overexpression resulted in an activation of ISGs, *VILMIR* may either act as a guide to recruit FUBP1 to enhance transcription, or *VILMIR* could act as a decoy to prevent FIR/PUF60 from negatively regulating transcription.

**Figure 5.**
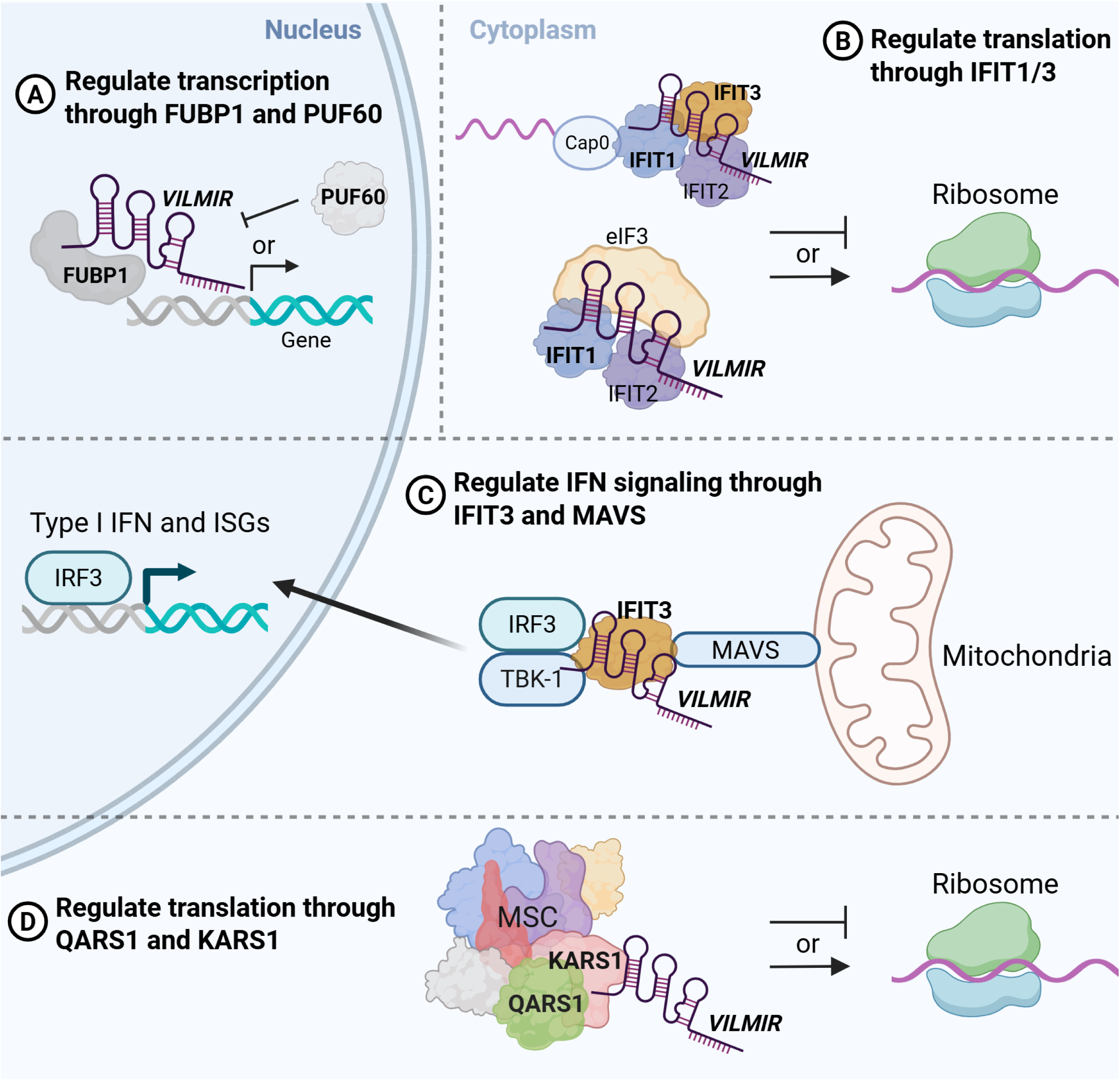
Schematic diagram of potential models of *VILMIR* function in the nucleus and cytoplasm to regulate host interferon responses. (A) *VILMIR* may interact with FUBP1 and PUF60 in the nucleus to regulate gene transcription, either by acting as a guide to recruit FUBP1 to enhance transcription or as a decoy to prevent PUF60/FIR from negatively regulating transcription. (B) In the cytoplasm, *VILMIR* may interact with IFIT1 and/or IFIT3 to regulate translation of cellular mRNA and viral RNA, either by the association of IFIT proteins with the 5’ end of RNAs or association with eukaryotic initiation factor 3 (eIF3). (C) Additionally, *VILMIR* may act as a scaffold to help bridge the mitochondrial antiviral signaling (MAVS) complex and the TNFR-associated factor family member-associated NF-κB activator-binding kinase 1 (TBK1), which leads to phosphorylation of interferon response factor 3 (IRF3) and induction of IFN-β and ISG expression. (D) Finally, *VILMIR* may interact with QARS1 and KARS1 in the multi-tRNA synthetase complex (MSC) to either positively or negatively regulate translation. (Figure created in https://BioRender.com)

In cytoplasmic lysate, we confirmed the interaction of *VILMIR* with IFIT1 and IFIT3 *in vitro*. IFIT1 and IFIT3 are known to inhibit translation of both cellular mRNA and viral RNA by either interacting with eukaryotic initiation factor 3 (eIF3) to block translation (31) or by binding directly to the 5’ end of non-self RNAs (32, 33). Therefore, we suggest a second model by which *VILMIR* either stabilizes the functions of IFIT1/3 to inhibit translation, or interferes with their function to enhance translation (Figure 5B). Apart from regulating translation, IFIT3 can also modulate interferon signaling by acting as a bridge between the mitochondrial antiviral signaling (MAVS) complex and the TNFR-associated factor family member-associated NF-κB activator-binding kinase 1 (TBK1), which leads to phosphorylation of interferon response factor 3 (IRF3) and induction of IFN-β and ISG expression (44). Therefore, we suggest a third model by which *VILMIR* acts as a scaffold to help bridge IFIT3 to MAVS and TBK1, thus enhancing ISG expression, as we observed that *VILMIR* overexpression resulted in an activation of ISGs (Figure 5C). Although the IFIT proteins often act in a complex (31), we did not identify IFIT2 or IFIT5 proteins in the mass spectrometry, so it is unknown whether this is due to the sensitivity of the assay or if *VILMIR* does not directly interact with these two proteins. In addition, as IFIT1 and IFIT3 were identified individually by mass spectrometry, the possibility that *VILMIR* interacts with these proteins in a complex needs to be further explored. While previous studies have reported lncRNAs that regulate the transcription of *IFIT* genes (45, 46), and another study reported that a segment of the lncRNA NORAD binds to IFIT proteins (47), to our knowledge, this is the first reported case of a full-length lncRNA interacting with IFIT proteins.

Finally, we also confirmed the interaction of *VILMIR* with two aminoacyl-tRNA synthetases (ARSs) *in-vitro*, QARS1 and KARS1. As stated above, ARSs are enzymes responsible for pairing tRNAs with amino acids during translation and also help form the multi-tRNA synthetase complex (MSC) (34). A study in 2020 found that mascRNA, a small RNA derived from the lncRNA *MALAT1*, binds to QARS1 in the MSC in order to promote global protein translation by regulating QARS1 protein levels (48). Similarly, we suggest a final model by which *VILMIR* either stabilizes QARS1 and/or KARS1 in the MSC to promote translation or interferes with their function to negatively regulate translation (Figure 5D). While these potential models need to be investigated further, they suggest that *VILMIR* may regulate host interferon responses through protein partners in both the nucleus and cytoplasm, which provides important foundational work in interrogating the specific mechanism of *VILMIR*.

As the pull-down assay was performed *in vitro,* future work is needed to confirm these protein interactions in cells, such as by an RNA immunoprecipitation (RIP) or RNA antisense purification (RAP) assay to establish their biological significance (49). Additional interacting molecules of *VILMIR* may also be determined by assays such as chromatin isolation by RNA purification (ChIRP), that can identify both chromatin and protein associations (49). This could also help determine if *VILMIR* is directly regulating the transcription of specific genes. In addition, immunoprecipitation assays could determine if *VILMIR* was binding to a unique protein or interacting with a protein complex. Finally, the biological significance of these protein interactions could be investigated by blocking or mutating the binding site on either *VILMIR* or the protein, such as in previous studies (14, 50).

In summary, we found that *VILMIR* interacts with multiple proteins in nuclear and cytoplasmic lysate. We also confirmed that *VILMIR* plays an activating role in the host interferon response in *trans* through the establishment of an overexpression cell line. Finally, by compiling RNA-seq analyses, we identified a core set of genes that are consistently differentially expressed after both *VILMIR* knockdown and overexpression. We have proposed several potential models for how *VILMIR* may function in the host interferon response. We expect these results will serve as a guide in probing the molecular mechanisms of *VILMIR* in detail, providing new insights into the biological significance of *VILMIR* during antiviral and interferon responses.

## ACKNOWLEDGMENTS

The authors would like to thank members of the Carpenter lab from the University of California, Santa Cruz, for their helpful advice in performing the RNA pull-down assay as well as establishing an overexpression cell line. In addition, Barbara Sherry, Kenneth Adler, Elisa Crisci, and members of the Peng lab from North Carolina State University College of Veterinary Medicine provided helpful feedback and critiques in manuscript preparation.

## DATA AVAILABILITY STATEMENT

The transcriptomic data discussed in this publication have been deposited in the Gene Expression Omnibus (GEO) https://www.ncbi.nlm.nih.gov/geo/ under accession numbers GSE261920 and GSE305270.

